# Population structure promotes the evolution of costly sex in artificial gene networks

**DOI:** 10.1101/333377

**Keywords:** Recombination, Hill-Robertson, Genetic architecture, Population structure, Cost of sex

## Abstract

We build on previous observations that Hill-Robertson interference generates an advantage of sex that, in structured populations, can be large enough to explain the evolutionary maintenance of costly sex. We employed a gene network model that explicitly incorporates interactions between genes. Mutations in the gene networks have variable effects that depend on the genetic background in which they appear. Consequently, our simulations include two costs of sex—recombination and migration loads—that were missing from previous studies of the evolution of costly sex. Our results suggest a critical role for population structure that lies in its ability to align the long- and short-term advantages of sex. We show that the addition of population structure favored the evolution of sex by disproportionately decreasing the equilibrium mean fitness of asexual populations, primarily by increasing the strength of Muller’s Ratchet. Population structure also increased the ability of the short-term advantage of sex to counter the primary limit to the evolution of sex in the gene network model—recombination load. On the other hand, highly structured populations experienced migration load in the form of Dobzhansky–Muller incompatibilities, decreasing the effective rate of migration between demes and, consequently, accelerating the accumulation of drift load in the *sexual* populations.

## Introduction

Sexual reproduction among eukaryotes is nearly ubiquitous (Bell 1982; Maynard Smith 1978; Vrijenhoek 1998) despite its substantial costs. Foremost among these is the risk that successful, coevolved genetic interactions will be destroyed by recombination. The production of males introduces a further cost, which may approach twofold in anisogamous species (Lehtonen et al. 2012; Gibson et al. 2017). Mating can also be a costly endeavor. In many species, an individual must expend time and resources to locate a mate, risking disease and predation in the process. The widespread prevalence of sex indicates it must convey extraordinary benefits, but the conditions that allow these benefits to counter its significant costs remain controversial.

One benefit of sex lies in its ability to increase the efficiency of selection by breaking up deleterious genetic associations (Burt 2000; Otto and Lenormand 2002). Such deleterious genetic associations can accumulate in finite populations through the combined activities of selection and drift, producing a phenomenon known as Hill-Robertson interference (Hill and Robertson 1966; Felsenstein 1974; Comeron et al. 2008). Hill-Robertson interference impairs evolvability and, in its strongest form, leads to the irreversible accumulation of deleterious mutations via a mechanism known as Muller’s Ratchet (Muller 1964; Felsenstein 1974; Haigh 1978). Recombination counters Hill-Robertson interference, generating an advantage of sex. Prior theoretical work on panmictic populations demonstrated that Hill-Robertson interference, particularly in the form of Muller’s Ratchet, facilitates the spread of modifiers that increase the recombination rate, that is, the *origin* of sex (Otto and Barton 2001; Iles et al. 2003; Barton and Otto 2005; Keightley and Otto 2006; Gordo and Campos 2008; Hartfield et al. 2010). However, the advantage of sex in pan-mictic populations was insufficient to allow the evolution of *costly* sex, except when the modifier led to only a small increase in the probability that sex occurs (Keightley and Otto 2006; Hartfield et al. 2010). The advantage of sex was stronger in structured populations, where costly sex was able both to evolve (Martin et al. 2006) and to resist invasion by asexual mutants (Peck et al. 1999; Salathé et al. 2006; Hartfield et al. 2012). This advantage of sex arising from population structure is expected to operate broadly because all natural populations show at least some structure.

The extent to which these findings can be generalized to real organisms, however, is unclear, as most of these models employed a simple genetic architecture, with limited to no interaction between genes (but see Peck et al. 1999). This precluded an important cost of sex—recombination load, which results from the disruption of coadapted gene complexes. This distinction proved vital in previous work that investigated the evolution of sex in an artificial gene network model that explicitly incorporated interactions between genes (Azevedo et al. 2006; Lohaus et al. 2010; Whitlock et al. 2016). As in real biological systems, mutations in the gene networks have variable effects that depend on the genetic background in which they appear. In panmictic populations of artificial gene networks, the interactions between genes evolved to produce a long-term (equilibrium fitness) advantage for sex that was strongest in small populations (*N <* 10^2^). By contrast, sex exhibited a short-term (invasibility) advantage only in the largest populations (*N >* 10^3^). Recombination load prevented both the origin and maintenance of sex in the small populations in which sex produced a large fitness advantage. As a result of the mismatch between the population sizes that produced the strongest short- and long-term advantages of sex, recombination load prevented the evolution of costly sex entirely (Whitlock et al. 2016).

We now ask whether population structure can facilitate the evolution of costly sex in the artificial gene network model. Population structure is expected to promote the evolution of sex because it introduces two processes that have a disproportionate negative impact on asexual populations. First, population structure strengthens the influence of Hill-Robertson interference (Martin et al. 2006; Hartfield et al. 2012). Second, it increases the time to fixation of invading asexual mutants, allowing more time for the operation of Hill-Robertson interference and, in particular, Muller’s Ratchet (Peck et al. 1999; Salathé et al. 2006). Because the short- and long-term advantages of sex depend, respectively, on the first (Gordo and Campos 2008) and second (Hartfield et al. 2012; Whitlock et al. 2016) of these effects, population structure has the potential to simultaneously increase both the short- and long-term advantages of sex, a possibility that has not been explored either theoretically or through simulation.

We note, however, that the complex genetic architecture of the gene network model, when combined with population structure, imposes another cost of sex: migration load. If we allow natural selection to operate in a spatially homogeneous manner, migrants will be equally well adapted to any deme, regardless of their sexual strategy. The offspring of sexual migrants will, nonetheless, experience a reduction in fitness (i.e., migration load) if they carry alleles that are incompatible with alleles in resident genotypes. Such Dobzhansky-Muller incompatilities (Dobzhansky 1937; Muller 1942) are common in nature, segregate within species (Corbett-Detig et al. 2013), and have been observed to evolve in a similar model (Palmer and Feldman 2009). Migration load can therefore be understood as an extension of recombination load, in which recombination between residents of different demes disrupts coadapted gene complexes and produces low-fitness hybrids. The impact of migration load on the benefits of sex imparted by population structure is unknown.

Here we evaluate the extent to which population structure supports the origin and maintenance of costly sex in a model with complex, evolving genetic architecture. We found that the relationship between population structure and the advantage of sex is non-monotonic. Small amounts of population structure did simultaneously increase both the short- and long-term advantages of sex by disproportionately increasing both the strength of Muller’s Ratchet in asexual populations and the time to fixation of invading asexual or sexual mutants. As a result, population structure enabled both the origin and maintenance of sex in the face of substantial costs. However, these advantages of sex were maximized in moderately structured populations. Highly structured sexual populations accumulated Dobzhansky–Muller incompatibilities that effectively prevented gene flow and, in some cases, disproportionately accelerated the accumulation of drift load (an analog of Muller’s Ratchet) in *sexual* populations.

## Methods

The gene network model used here is based on a model introduced by Wagner (1994,1996).

### Genotype

A haploid genotype is modeled as a network of *n* genes, each encoding a transcription factor that can, potentially, regulate its own expression or the expression of other genes. The gene network is represented by an *n × n* matrix, **R**, where *r_ij_* ∈ **R** is the regulatory effect of the product of gene *j* on gene *i*.

### Phenotype

The expression pattern of an individual is represented by the vector **S**, where *s_i_* ∈ **S** is the expression state of gene *i* = 1, 2, …, *n*. Expression states are discrete: a gene is either on (*s_i_* = +1) or off (*s_i_* = *−*1).

The expression pattern of an individual at time *t* is given by the system of difference equations

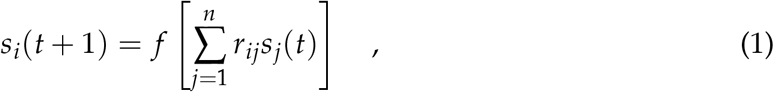

where *f* is a step function that determines how the input from the gene network controls the expression of the target gene:

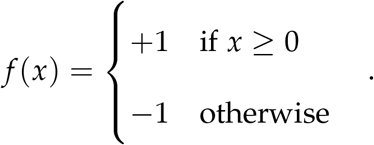

Starting from an initial expression pattern **S**(0) at time *t* = 0, gene expression changes according to Equation 1 and is judged to reach a steady state if the following criterion is met: **S**(*t*) = **S**(*t −* 1). If a genotype does not achieve a gene expression steady state within *t ≤* 100 time steps, it is considered inviable (*W* = 0, see next section). If a genotype achieves a gene expression steady state within *t ≤* 100 time steps, it is considered viable (*W >* 0), and the steady state gene expression pattern **Ŝ** is its *phenotype*.

### Fitness

The fitness of a viable genotype is given by

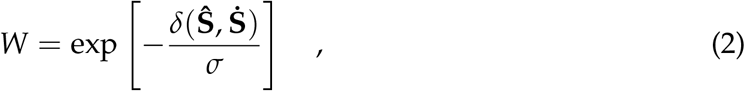

where 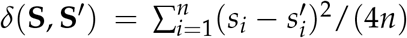 measures the difference between expression patterns **S** and **S′**, **Ŝ** is the phenotype corresponding to the genotype, **Ṡ** is the optimal phenotype, and *σ >* 0 is inversely related to the strength of stabilizing selection.

Offspring of asexual reproduction have fitness *W*_asex_ = *W*, whereas offspring of sexual reproduction have fitness *W*_sex_ = *W*/*C*, where *C ≥* 1 is the cost of sex. For example, *C* = 1 and *C* = 2 represent no cost and a two-fold cost, respectively.

### Random genotype

A random genotype is created by generating a random gene network and a random initial gene expression pattern. A random gene network is obtained by randomly filling an **R** matrix with (1 *− γ*)*n*^2^ zeros and *γn*^2^ random numbers from a standard normal distribution, where *γ* is the connectivity density of the network. A random initial gene expression pattern is generated by filling the *n* entries of **S**(0) with either *−*1 or +1 with equal probability.

### Evolution

Evolution is simulated using an individual-based, Wright-Fisher model with non-overlapping generations. Populations maintain a constant total size, *N*, and are subdivided into *D* demes of equal size, *N_d_* = *N*/*D*, arranged in a ring. In each generation, individuals undergo a selection-reproduction-mutation-migration life cycle.

#### Initialization

Simulations are initiated by producing *N* clones of a single viable randomly generated genotype. The phenotype of this founding individual is set as the population’s optimal phenotype. The optimum remains constant over the course of the simulation.

#### Selection

Every generation, in each deme, the individuals that will reproduce are chosen at random, with replacement, with probability proportional to their fitness (Equation 2).

#### Reproduction

If the parent reproduces asexually, it generates a clone of itself; if a parent reproduces sexually, it mates with another sexually reproducing parent and produces one haploid recombinant offspring. The recombinant **R** matrix is generated by copying rows from the **R** matrices of the parents with equal probability. This is equivalent to free recombination between regulatory regions and no recombination within regulatory regions.

#### Mutation

Each individual offspring acquires a random number of mutations drawn from a Poisson distribution with mean *U* (the genomic mutation rate). A mutation is represented by a change to the value of one of the *γn*^2^ nonzero entries in **R** chosen at random; the mutated value is drawn randomly from a standard normal distribution.

#### Migration

Every generation, a number of individuals from each deme is randomly chosen (without replacement) to migrate to either neighboring deme, with no bias for fitness or reproductive mode. The number of migrants from one deme to one of its neighbors is Poisson distributed with parameter *mN_d_*/2, where *m* is the migration rate. The numbers of migrants to and from individual demes are not guaranteed to be equal. Thus, following migration the population may transiently show variation in deme sizes. Selection and reproduction restore the size of every deme to *N_d_*.

### Reproductive mode

The reproductive mode of an individual is determined by its genotype at a modifier locus, *rec*, unlinked to the genes involved in the gene network. There are two alleles at the modifier locus: *rec^−^* and *rec^+^*. If a population is fixed for the *rec^−^* allele, every individual reproduces asexually, and if it is fixed for the *rec^+^* allele, every individual reproduces sexually.

The sexual and asexual subpopulations are reproductively isolated from each other. Sexual organisms do not experience a frequency-dependent cost of finding mates. One individual is chosen for every reproductive event with probability proportional to its fitness. If it carries the *rec^−^* allele, it reproduces asexually. If it carries the *rec^+^* allele, a second individual carrying an *rec^+^* allele is chosen with probability proportional to its fitness, and the two individuals reproduce sexually and produce one recombinant offspring. The second individual may be the same as the first individual because sampling is done with replacement; the resulting reproductive event is equivalent to asexual reproduction.

### Invasion analyses

Populations were evolved for 10^4^ generations under asexual (or sexual) reproduction to allow sufficient time for the population to approach mutation-recombination-selection-drift-migration equilibrium. We then mutated the allele at the modifier locus *rec* (*rec^−^ ↔ rec^+^*, see “Reproductive mode” above) in a single randomly chosen individual, causing it to switch to the opposite reproductive mode, from asexual to sexual or vice-versa. We measured the fixation probability of the novel modifier allele, *u*, relative to that of a neutral mutation (*u^*^* = 1/*N*) in replicate invasion trials. During the invasion trials, individuals carrying the sexual modifier allele *rec^+^* experience a fixed cost of sex, *C* (see “Fitness” above). For invasion trials in which the novel modifier allele fixed, we recorded the generation in which it fixed as *T*_fix_.

### Population metrics

#### Mean fitness

The mean fitness, *W̅*, of all individuals present in the population at a given time (see Equation 2). The mean fitness of sexual and asexual individuals is denoted by *W̅*_sex_ and *W̅*_asex_, respectively.

#### Rate of adaptation

Defined as

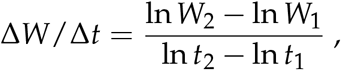

where *W_i_* is mean fitness at generation *t_i_*. We used *t*_1_ = 562 and *t*_2_ = 1778 generations in our calculations because variance among treatments in ∆*W*/∆*t* was maximized over this time interval.

#### Deleterious mutation rate

Defined as *U_d_* = *U*(*p_l_* + *p_d_*), where *U* is the genomic mutation rate and *p_l_* and *p_d_* are the proportion of lethal and non-lethal deleterious mutations, respectively. *U* is constant throughout the course of a simulation but *p_l_* and *p_d_* can evolve. We estimated the quantity *p_l_* + *p_d_* for a genotype by generating 100 copies of the genotype, each carrying one independently generated mutation, and evaluating the proportion of them that have lower fitness than the original genotype.

#### Epistasis

We define multiplicative epistasis between two nonlethal mutations, *i* and *j*, as *ε* = *W_ij_*/*W − W_i_W_j_*/*W*^2^, where *W* is the fitness of the unmutated genotype, *W_i_* and *W_j_* are the fitnesses of the single mutants, and *W_ij_* is the fitness of the double mutant. We calculated mean epistasis coefficients, *ε*, across a random sample of 100 pairs of mutations, introduced individually and in combination into each viable individual in the population.

#### Mean fitness of recombinant offspring

Defined as *W̅_R_*, the mean fitness of offspring produced by recombination (without mutation) between individuals of the same deme, *W_r_*, relative to the average fitness of its parents *W̅_p_*. We averaged *W_r_*/*W̅_p_* across *N* independently chosen pairs of individuals, where each parent was chosen with probability proportional to its fitness (i.e., in the same way the population reproduced in the evolutionary simulations).

#### Mean fitness of migrant offspring

Defined as *W̅_M_*, the mean fitness of offspring produced by recombination (without mutation) between individuals of neighboring demes, *W_m_*, relative to the average fitness of its parents *W̅_p_*. We averaged *W_m_*/*W̅_p_* across *N* independent offspring, generated by recombination between *N_d_* pairs of parents randomly chosen from each of the *D* pairs of neighboring demes. Both parents were chosen from among the individuals in their respective demes with probability proportional their fitness.

#### Population differentiation

Measured as *Q_ST_* = *V_b_*/(*V_w_* + *V_b_*), where *V_w_* and *V_b_* are the variance within and among demes at a neutral locus, respectively. Each individual carries 20 neutral loci, which were neither linked to each other nor to any of the gene network loci. The founding genotype had a value of 0 at all 20 neutral loci. Every generation, each neutral locus in each individual acquired a random number of mutations drawn from a Poisson distribution with mean 1.0 (the neutral locus mutation rate). Each mutation changed the value of the neutral locus by adding a random value drawn from the standard normal distribution. *V_w_* and *V_b_* were calculated as variance components in a random-effects analysis of variance. We averaged *Q_ST_* across the 20 neutral loci.

#### Maximum cost of sex

Defined as origin-*C*_max_ and maintenance-*C*_max_ to describe the maximum sustainable cost of sex during the origin and maintenance of sex, respectively. Origin-*C*_max_ is the maximum value of *C* for which *u*_sex_ *> u^*^* in sexual invasion trials. Maintenance-*C*_max_ is the maximum value of *C* for which *u*_asex_ *< u^*^* in asexual invasion trials (see “Invasion analyses”). For each set of equilibrium populations evolved at a particular *m* and *N_d_*, we measured the modifier fixation probability, *u*, at each of at least 4 values of *C* and estimated *C*_max_ from the resulting data using logistic regression (Figures S2 and S3).

#### Equilibrium values

Defined as values of population metrics at generation 104 and in-dicated by a hat symbol (e.g., *Ŵ*sex).

### Parameter values

By default, we used the same parameters as in Whitlock et al. (2016): networks of *n* = 100 genes with connectivity density *c* = 0.05, a genomic mutation rate of *U* = 1 per generation, and moderate stabilizing selection (*σ* = 0.2). Using these parameters, mutations have a broad range of potential fitness effects, including beneficial, neutral, slightly deleterious and lethal (Figure S3). Random genotypes have a deleterious mutation rate of *U_d_ ≈* 0.5 (Figure S2, Whitlock et al. 2016). This value is comparable to those estimated for fruitflies (*U_d_* = 0.6, Haag-Liautard et al. 2008) and rodents (*U_d_* = 0.5, Gaffney and Keightley 2006).

We simulate populations of *N* = 10^3^ individuals. We chose this value because it is the largest population size in which sex is approximately neutral in the short term in panmictic populations (see Fig. 4 of Whitlock et al. 2016).

### Statistical Analysis

All statistics were conducted using the R statistical package, version 3.4.1 (Ihaka and Gentleman 1996). Logistic regression was conducted using the function *glm* from the *stats* package. Linear models that investigated the contributions of various population metrics to the long- and short-term advantages of sex were conducted using the function *lm* from the *stats* package.

All data were generated from 50 pairs of sexual and asexual populations. Each pair was founded by an independent, randomly generated genotype. Combined metrics from sexual and asexual populations, like *Ŵ*_sex_/*Ŵ*_asex_, were calculated separately for each pair, and the resulting 50 composite values were used for plotting and statistical tests.

### Data Availability

Programs used to run all simulations were written in Python 2.7 and are available at https://bitbucket.org/aobwhitlock/whitlock-et-al-2018. Supplemental figures are available in the Supporting Information.

## Results

### Population structure has a complex effect on the long-term advantage of sex

Previously, it was shown that panmictic (i.e., unstructured) sexual populations of *N* = 10^3^ individuals experiencing a genomic mutation rate of *U* = 1 and moderate stabilizing selection (*σ* = 0.2) evolved a mean fitness at equilibrium 5.7% *±* 0.8% (mean and 95% confidence interval) higher than that of otherwise identical asexual populations (Whitlock et al. 2016). Here we investigate the extent to which population structure increases the long-term advantage of sex.

We manipulated population structure in two ways. First, we subdivided the total population into *D* = 10 to 100 demes of identical size such that *N_d_* = 10^3^/*D* (note that *D* and *N_d_* were not varied independently). Demes were arranged around a circle and migration occurred via a one-dimensional stepping-stone model (Kimura 1952). Second, we varied the migration rate (*m*) between neighboring demes from 2 *×* 10^−4^ to 0.2. There was no migration (i.e., *m* = 0) between non-neighboring demes. Structured populations consisting entirely of either sexual or asexual individuals were allowed to evolve for 10^4^ generations, by which time most populations had reached an approximate mutation-selection-drift-migration fitness equilibrium (Figure S1). The least structured populations (i.e., those subdivided into *D* = 10 demes of *N_d_* = 100 individuals each, experiencing the highest migration rate, *m* = 0.2) evolved approximately like panmictic populations, achieving similar equilibrium mean fitness and *Q_ST_ ≈* 0 (Figure 1). Decreasing either the deme size (*N_d_*) or the migration rate (*m*) caused the level of genetic differentiation among demes, *Q_ST_*, to increase, confirming that populations became more structured (Figure 1).

**Figure 1:**
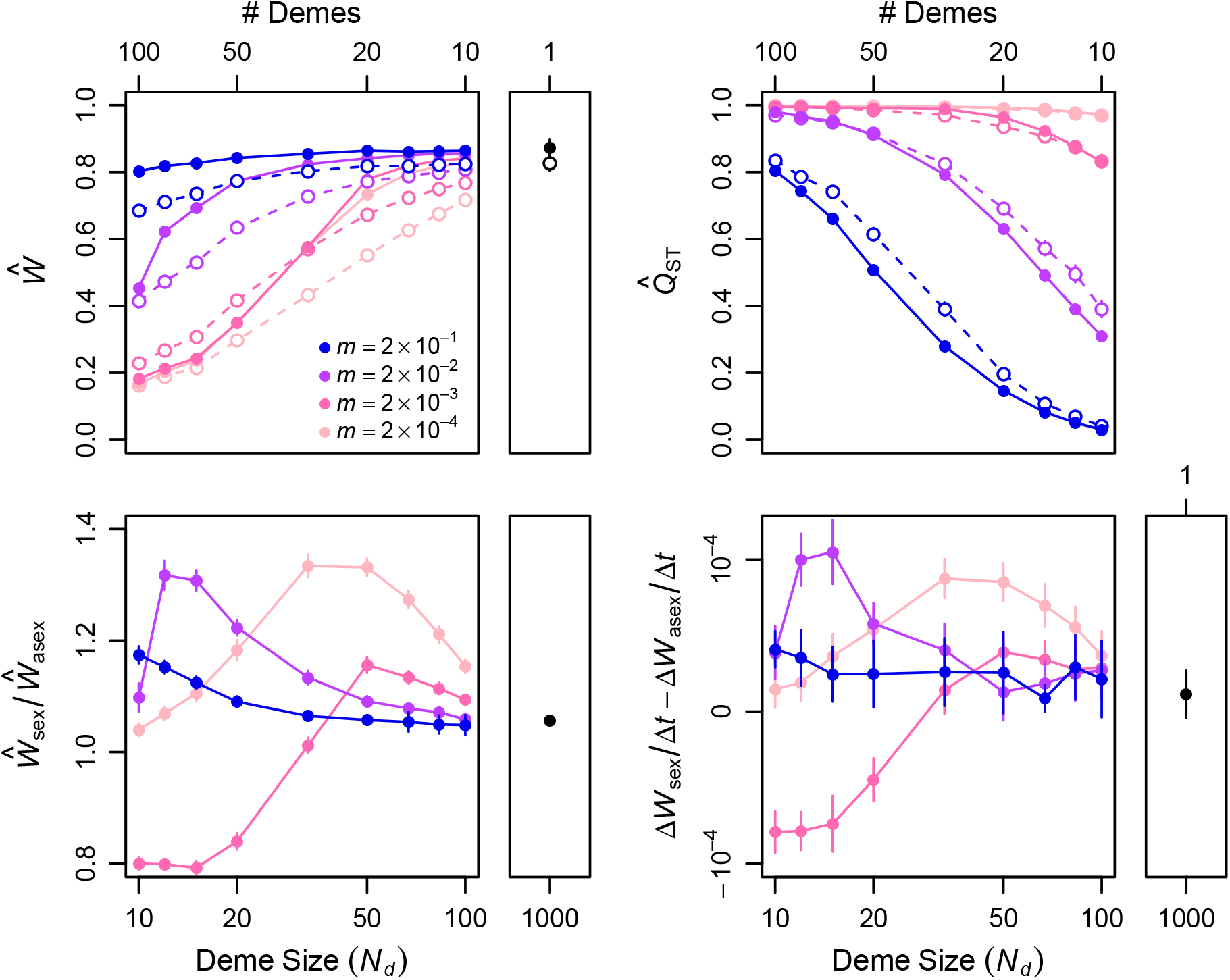
Migration rate and population subdivision interact in a complex way to determine the long-term advantage of sex. Mean fitness (*Ŵ*) the fitness advantage of sexuals (*Ŵ*_sex_/ *Ŵ*_asex_), and population differentiation (*Q̂_ST_*) were calculated at generation 10, after populations of all migration rates and deme sizes had achieved approximate fitness equilibria. The rate of adaptation relative (∆*W*_sex_/∆*t* ∆*W*_asex_/∆*t*) was calculated at generation 10^3^. In the top panels, sexual and asexual populations are represented by closed and open circles, respectively. Large panels show the relationships between these equilibrium metrics and the deme number and size of subdivided populations. Small panels show the equilibrium metrics calculated from panmictic asexual and sexual populations of the same total size, *N* = 10^3^ (obtained as in Whitlock et al. 2016). Values are means and 95% confidence intervals based on 50 replicate populations initiated from different randomly chosen founders.

Relative to panmictic populations, small increases in population structure, imposed by decreasing either *N_d_* or *m*, caused the magnitude of the long-term advantage of sex (*Ŵ*_sex_/*Ŵ*_asex_) to increase (Figure 1). This pattern was true provided the population consisted of relatively large demes (*N_d_* ≳ 50). At *N_d_* = 50 and *m* = 2 *×* 10^−4^, the average advantage of sex achieved a maximum that was 5.8-fold larger than it was in a panmictic population of the same total size.

Indeed, for each *m ≤* 2 *×* 10^−2^, there was an intermediate deme size, 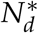, at which the advantage of sex was maximized. Further subdivision into demes that were smaller than 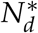 eroded the advantage of sex. Surprisingly, the effect of small deme size was most pronounced at the intermediate migration rate of *m* = 2 *×* 10^−3^, generating a strong *disadvantage* of sex in the smallest deme sizes (*N_d_ ≤* 33) at this migration rate.

The variance we observed in the long-term advantage of sex was almost entirely explained by the relative rate of adaptation in sexual compared to asexual populations (∆*W*_sex_/∆*t −* ∆*W*_asex_/∆*t*; *R*^2^ = 0.95, *F*_1,34_ = 669.3, *p <* 10^−15^; Figure 1). We note that ∆*W*/∆*t* is a composite metric that increases with adaptation rate and decreases with the accumulation of genetic load. Although ∆*W*/∆*t*, by definition, measures the rate of adaptation, in our experiments differences in ∆*W*/∆*t* may better reflect differences in the rate of accumulation of load. ∆*W*_sex_/∆*t −* ∆*W*_asex_/∆*t* took on its highest values for combinations of *m* and *N_d_* in which migration effectively countered the accumulation of drift load in sexual (∆*W*_sex_/∆*t >* 0) but not asexual (∆*W*_asex_/∆*t <* 0) populations. In these conditions, asexual populations experienced Muller’s Ratchet. Conditions in which drift load accumulated in both sexuals and asexuals, but did so more rapidly in sexuals (∆*W*_sex_/∆*t <* ∆*W*_asex_/∆*t <* 0), were entirely unexpected because Muller’s Ratchet is not expected to operate in sexual populations. We next investigated the cause of the accelerated accumulation of drift load in some of the highly structured sexual populations.

### Dobzhansky-Muller incompatibilities accelerate the accumulation of drift load in structured sexual populations

We hypothesized that evolving properties of the genetic architecture made substantial contributions to rate at which drift load accumulated, thus we measured the mutation, recombination, and migration loads of equilibrium structured populations. Both the mutation and recombination loads evolved in ways that contributed to the long-term advantage of sex (Figure 2), as was observed previously in panmictic populations (Whitlock et al. 2016). The lower equilibrium *Û_d_* of sexual, compared to asexual, populations generated a fitness advantage for sexual populations of 6–9% (1.06 *<* exp(*−Û_d_*_,sex_)/ exp(*−Û_d_*_,asex_) *<* 1.09). Recombination load at equilibrium generated a fitness disadvantage for sexual populations of 5–8% (0.92 *< Ŵ_R_<* 0.95). However, the differences among treatments in mutation and recombination loads (Figure 2) cannot explain the differences among treatments in the overall advantage to sex (Figure 1; linear regression of *Ŵ* sex/ *Ŵ*asex against exp(*−Ûd*,sex)/ exp(*–U Ŵ_d_*_,asex_): *R*^2^ = 0.0937, *F*_1,34_ = 3.514, *p* = 0.0695; linear regression against *ŴR*: *R*^2^ = 0.0153, *F*_1,34_ = 0.5275, *p* = 0.4726).

**Figure 2:**
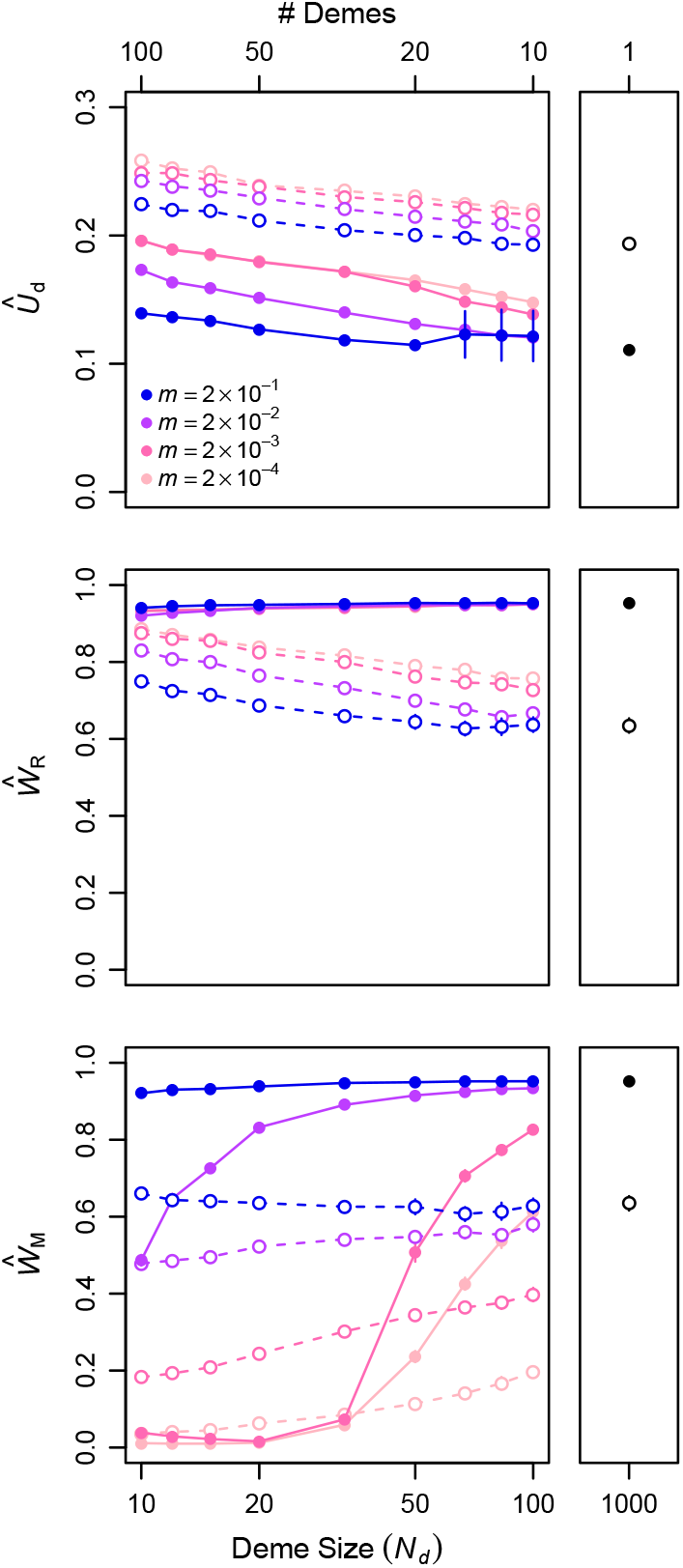
Migration rate and deme size affect three aspects of the equilibrium genetic architecture. Deleterious mutation rate (*Û_d_*), the mean fitness of recombinant offspring of parents from the same deme (*Ŵ_R_*), and the mean fitness of recombinant offspring of parents from different demes (*Ŵ_M_*) at generation 10^4^, after populations of all migration rates and deme sizes achieved approximate fitness equilibria. As in Figure 1, sexual and asexual populations are represented by closed and open circles, respectively, and values are means and 95% confidence intervals based on 50 replicate populations initiated from different randomly chosen founders. Large panels show the relationships between these equilibrium metrics and the deme number and size of subdivided populations. Small panels show the equilibrium metrics calculated from panmictic asexual and sexual populations of size *N* = 10^3^ (obtained as in Whitlock et al. 2016). Note that measures of *Ŵ_R_* and *Ŵ_M_* in asexual populations are provided for comparison only. During the evolution experiments, offspring produced by asexual reproduction experience *W_R_* = 1 and *W_M_* = 1, by definition.

A third property of the genetic architecture, the relative fitness of sexual migrant off-spring, *Ŵ_M_*, also evolved in our simulations (Figure 2). At equilibrium, *Ŵ_M_* depended predictably on both *N_d_* and *m*. For large *N_d_* and *m*, the fitness of migrant offspring was indistinguishable from that of resident offspring (i.e., *Ŵ_M_ ≈ Ŵ_R_*), as expected when there is relatively little differentiation between demes (low *Q̂_ST_*). As *N_d_* and *m* declined, so did *Ŵ_M_*, until when *N_d_* ≲ S 50 and *m ≤* 2 *×* 10^−3^, demes were strongly differentiated (high *Q̂_ST_*) and *Ŵ_M_ ≈* 0. These reductions in *Ŵ_M_* must have resulted from the accumulation of alleles in some demes that show strong negative epistatic interactions with alleles present in other demes—i.e., Dobzhansky-Muller incompatilities. This extreme variation in *Ŵ_M_* had only a negligible direct effect on population mean fitness because *Ŵ_M_* was low only when migration rates were also low (*m* ≲ S 2 *×* 10^−3^). Instead, the main consequence of a low *Ŵ_M_* was to reduce the effective gene flow between demes, which in turn intensified the accumulation of drift load in sexual populations.

In highly structured populations with moderate migration rates (e.g., *N_d_ <* 50 and *m* = 2 *×* 10^−3^; hot pink line in Figure 2), the accumulation of Dobzhansky-Muller incompatibilities (*W_M_ ≈* 0) explains the faster fitness declines in sexual than in asexual populations (∆*W*_sex_/∆*t <* ∆*W*_asex_/∆*t*; hot pink line in Figure 1). In this parameter range, the moderate migration experienced only by asexual populations more effectively countered Muller’s Ratchet than the recombination experienced only by sexual populations. Outside this parameter range, the effective migration rate was similar in asexual and sexual populations, either because Dobzhansky-Muller incompatibilities didn’t accumulate in sexual populations (*m >* 10^−3^) or because the migration rate was also near zero in asexual populations (*m* = 10^−4^).

### The long-term advantage of sex is a good predictor of the evolutionary success of sex

Previously, it was shown that sexual modifier mutations were approximately neutral in panmictic asexual populations of *N* = 10^3^; asexual modifier mutations were also approximately neutral in panmictic sexual populations of the same size. Thus, despite the existence of a long-term advantage of sex, sexual reproduction neither evolved nor was maintained in panmictic populations of *N* = 10^3^ individuals (Whitlock et al. 2016). The addition of population structure is expected to promote the maintenance of costly sex — perhaps even costly sex— by increasing the time (*T*_fix_) that asexual modifier mutations spend in transit to fixation. In turn, increasing *T*_fix_ is expected to increase the accumulation of deleterious mutations in the invading asexual lineage, depressing its mean fitness and probability of fixation. This effect, of population structure on *T*_fix_, is not expected to influence the *origin* of costly sex. Here we investigate the extent to which the predicted effect of population structure on *T*_fix_ combines with the complex effect of population structure on equilibrium mean fitness (Figure 1) to promote both the origin and maintenance of costly sex.

As expected, population structure promoted the maintenance of costly sex (Figures 3B and 3D). For each combination of *m* and *N_d_* we used invasion assays to estimate the maximum cost of sex (maintenance-*C*_max_) under which the fixation probability of an asexual modifier mutation was lower than the neutral expectation (*u < u^*^*, Figure S2). maintenance-*C*_max_ *>* 1 for almost all the parameter combinations we examined and rose as high as 1.36 for the most favorable parameter combination. Maintenance-*C*_max_ significantly increased with both the mean *T*_fix_ among successful invaders (*F*_1,13_ = 142.5, *p <* 10^−7^) and the equilibrium advantage to sex (*Ŵ*_sex_/*Ŵ*_asex_, *F*_1,13_ = 343.7, *p <* 10^−10^). Indeed, *T*_fix_ and the equilibrium advantage of sex predicted maintenance-*C*_max_ almost perfectly (multiple regression model containing only these two predictors, *R*^2^ = 0.98).

**Figure 3:**
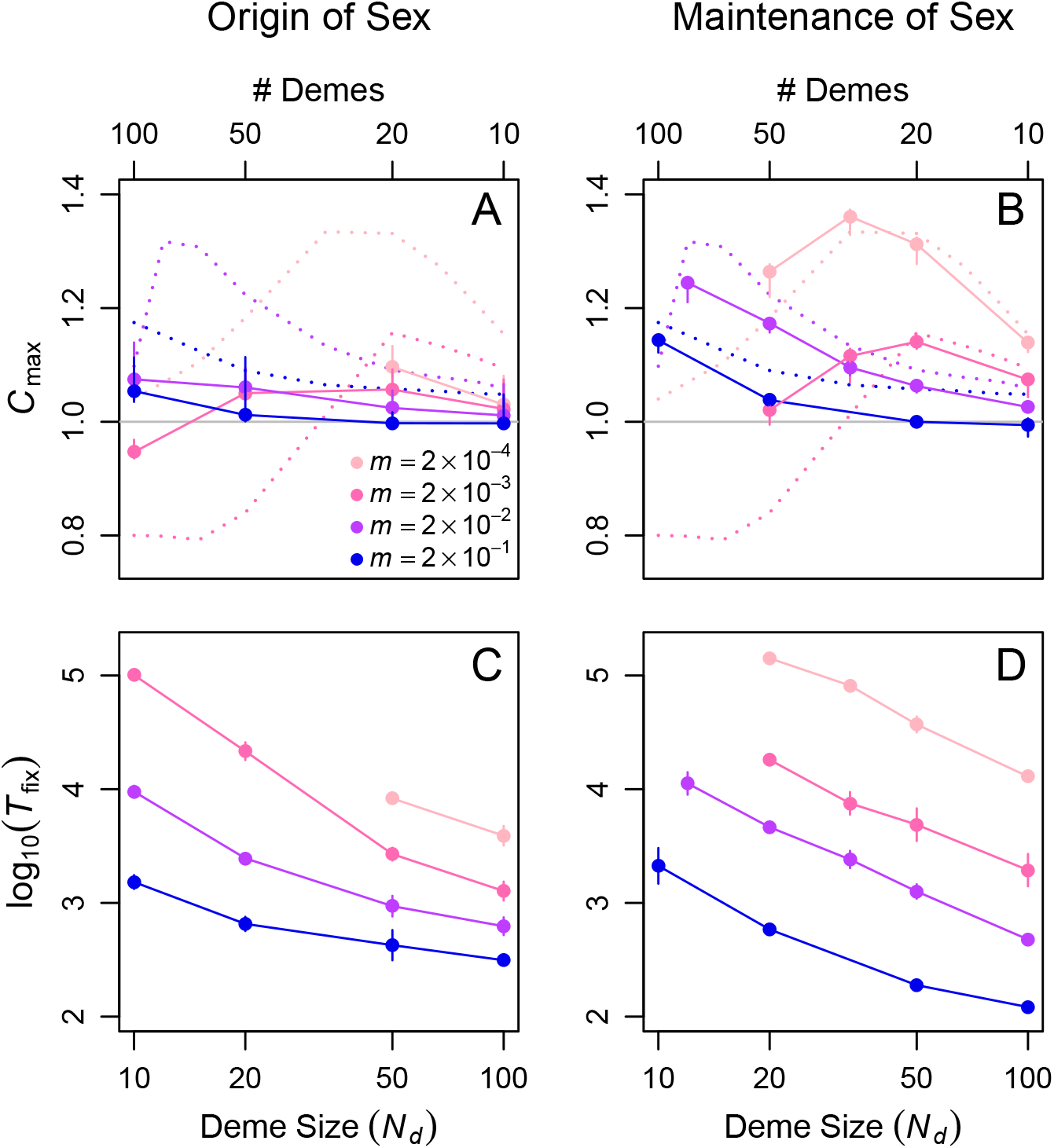
Population structure increases the transit time (*T*_fix_) of invading sexual and asexual mutants, facilitating both the origin and maintenance of costly sex. (A) Points connected by solid lines show the maximum cost of sex (*C*_max_) at which sexual mutants successfully invaded equilibrium asexual populations in each deme size and migration rate treatment. (B) Points connected by solid lines show the maximum cost of sex at which equilibrium sexual populations successfully resisted invasion by asexual mutants. Dotted lines in both (A) and (B) show the long-term advantage of sex, *Ŵ*_sex_/ *Ŵ*_asex_, replotted from Figure 1, for comparison. (C and D) The average fixation time (*T*_fix_) of sexual (C) and asexual (D) mutants that successfully invaded equilibrium asexual and sexual populations, respectively. Values are means and 95% confidence intervals based on all of the successful invasions in each condition. Missing points in the *m* = 2 × 10^−3^ and *m* = 2 × 10^−4^ series correspond to parameter combinations for which invasion trials were not run because they were not computationally feasible (i.e., *T*_fix_ was too long).

Population structure also promoted the origin of costly sex (Figures 3A and 3C). To investigate the origin of sex we used invasion assays to estimate the maximum cost of sex (origin-*C*_max_) under which the fixation probability of a sexual modifier mutation was higher than the neutral expectation (*u > u^*^*, Figure S3). Similar to maintenance-*C*_max_, origin-*C*_max_ *>* 1 for almost all the parameter combinations we examined and increased significantly with the equilibrium advantage to sex (*F*_1,11_ = 12.71, *p* = 0.0044). However, origin-*C*_max_ rose only as high as 1.1 for the most favorable parameter combination and showed no significant relationship with *T*_fix_ (*F*_1,11_ = 3.62, *p* = 0.0835). In addition, origin-*C*_max_ was only somewhat predictable (*R*^2^ = 0.60) from the quantities *Ŵ* sex/*Ŵ*asex and *T*_fix_.

We note that these results are for modifiers of sex that cause free recombination between rows of the genotype matrix, **R**. We also examined the behavior of modifiers of recombination that cause only small changes in recombination rate, using a model that allows recurrent mutation at the modifier locus. Modifiers of recombination behaved similarly to modifiers of sex. Using the parameter combination that was most favorable to the origin of sex, recombination rate increased over time when *C* = 1.1, but not when *C* = 1.2 (Figure 4). This similarity between the two types of modifiers was not observed in panmictic populations, where modifiers of recombination were more likely to evolve than modifiers of sex (Whitlock et al. 2016).

**Figure 4:**
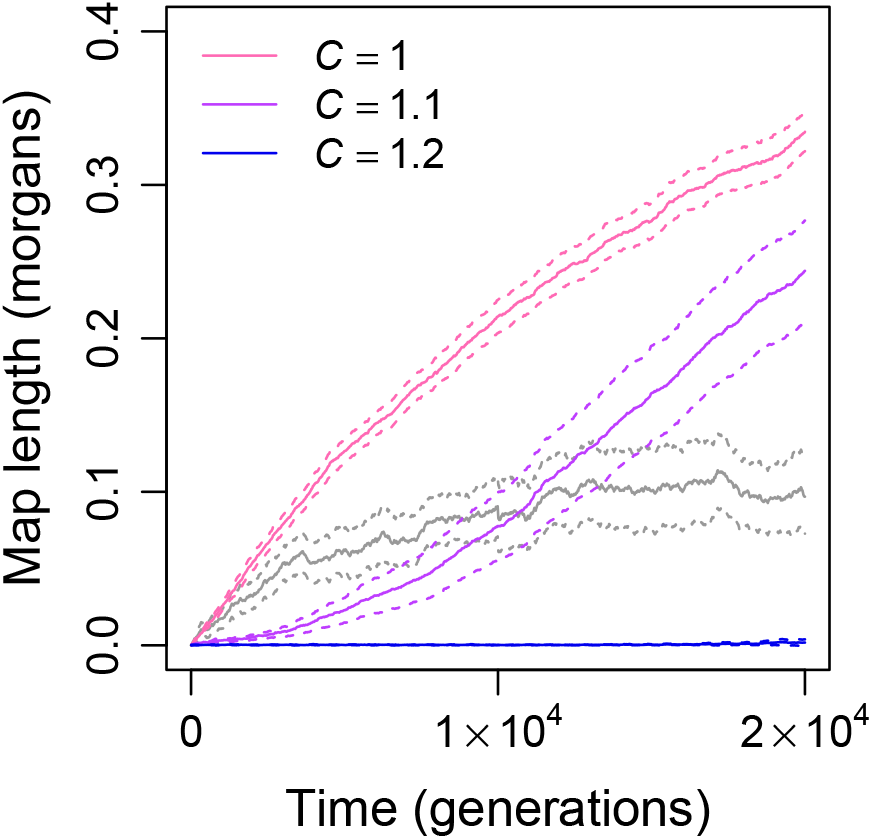
Evolution of the recombination rate under recurrent mutation at the recombination modifier locus. Color lines show the change in mean and 95% confidence interval, respectively, of the genome map length (i.e., mean crossover probability) over time in structured populations with *N_d_* = 50 and *m* = 2 × 10^−4^ and costs of sex ranging from none (*C* = 1, pink) to substantial (purple and blue). Panmictic populations of the same total size (*N* = 10^3^) experiencing no cost of sex (*C* = 1) are shown in gray for comparison. Data are from 50 replicates initiated with the equilibrium structured asexual populations shown in Figures 1 and S1 (pink, purple, and blue) or the equilibrium panmictic asexual populations of size *N* = 10^3^ (gray, as in Whitlock et al. 2016).

## Discussion

Our work builds on prior observations that Hill-Robertson interference generates an advantage of sex that, in structured populations, can be large enough to explain the evolutionary maintenance of *costly* sex (Peck et al. 1999; Salathé et al. 2006; Martin and Lenormand 2006; Hartfield et al. 2012). We employed an artificial gene network model that explicitly includes genetic interactions, resulting in two limits on the evolution of costly sex that were not present in the prior work—recombination load and migration load. First, population structure improved the ability of the short-term advantage of sex to counter recombination load. As a result, population structure facilitated the *origin* of sex, in addition to its maintenance. Second, highly structured populations experienced migration load in the form of Dobzhansky-Muller incompatibilities, accelerating the accumulation of drift load in the most-structured sexual populations and reducing the long-term advantage of sex. This effect was responsible for the observation that the maximum advantage of sex was realized in moderately structured populations.

Our first observation, that structure promoted the origin of sex, was not predictable from prior investigations of the evolution of sex because it resulted from the complex genetic architecture in the gene network model. Sexual lineages experience a short-term fitness decline, in the form of recombination load, before they evolve robustness that ameliorates the accumulating recombination load and produces a long-term fitness advantage in the form of lower *U_d_*. (Figures S1, 1, and 2). In panmictic populations, successful sexual invaders must arise by chance in a high fitness genetic background and quickly hitchhike with that genetic background to a frequency that prevents their loss during the short-term fitness decline (Whitlock et al. 2016). Population structure facilitated the origin of sex by slowing the loss of such sexual invaders during the short-term fitness decline. In other words, population structure reduced the limitation imposed by recombination load by increasing the time over which recombinational robustness can evolve (*T_f ix_*) in invading sexual lineages.

Our second observation, that the long-term advantage of sex was maximized at intermediate structure, was also not predictable from prior work conducted using simpler genetic architectures. In the gene network model, differences in the rate at which drift load accumulated in sexual and asexual populations depended, in turn, on the effective rate of migration between demes, *m_e_ ≈ m W̅_M_*. Increasing structure promoted the accumulation of Dobzhansky-Muller incompatibilities in sexual populations (Dobzhansky 1937; Muller 1942; Palmer and Feldman 2009; Bank et al. 2012), which in turn decreased the effective migration rate and resulted in substantial increases in migration load. In the most structured sexual populations, the mean fitness of hybrids between demes dropped to *W̅_M_ ≈* 0 and the effective migration rate dropped to *m_e_ ≈* 0. Asexual populations did not experience migration load (i.e., *W̅_M_* = 1 and *m_e_* = *m*) because asexual migrant alleles did not change genetic backgrounds and were equally well-adapted to local and distant environments. Thus, asexual populations benefited from migration in some high-structure conditions (e.g., *m* = 10^−3^ and *N_d_ <* 33) where sexual populations did not. In these conditions, drift load actually accumulated faster in sexual than in asexual populations. Strong Dobzhansky-Muller genetic incompatilities are observed in other similar models (Palmer and Feldman 2009), and both within (Corbett-Detig et al. 2013) and among (Presgraves 2010; Maheshwari and Barbash 2011) biological species.

Earlier work has shown that migration load could generate a cost of sex in structured populations experiencing spatially *heterogeneous* selection (García-Ramos and Kirkpatrick 1997; Kirkpatrick and Barton 1997; Hendry et al. 2001; Alleaume-Benharira et al. 2006; Bolnick and Nosil 2007). Our results extend the potential role of migration load as a cost of sex to structured populations under spatially *homogeneous* selection. Our data also revealed a general mechanism for the cost of sex caused by migration load. Migration load, whether generated by the accumulation of Dobzhansky-Muller incompatibilities or by local maladaptation, reduces the effective migration rate, which intensifies the accumulation of drift load in sexual populations.

In sum, our results demonstrate that complex genetic architecture interacts with population structure to both promote and constrain the origin and maintenance of costly sex. The magnitude of both the long- and short-term advantages of sex are likely to be affected by many factors not considered here, including deviations from random mating other than those arising from population structure, ploidy, and environmental change (Shields 1982; Agrawal 2001; Siller 2001; Blachford and Agrawal 2006; Kirkpatrick and Jenkins 1989; Kondrashov and Crow 1991; Agrawal and Chasnov 2001; Otto 2003; Charlesworth 1993; Barton 1995; Otto and Nuismer 2004; Carja et al. 2014; Nowak et al. 2014). We posit that these factors may also interact with a complex genetic architecture to influence the evolution of sex, making them promising avenues for future inquiry.

## Acknowledgements

We thank Sonia Singhal for her comments on the manuscript. The University of North Carolina at Chapel Hill and the Research Computing group provided computational resources and support. This work was funded by grant DEB-1355084 awarded to C.L.B. and grant DEB-1354952 awarded to R.B.R.A. by the National Science Foundation and by a Dissertation Completion Fellowship awarded to A.O.B.W. by the University of North Carolina at Chapel Hill. The funders had no role in study design, data collection and analysis, decision to publish, or preparation of the manuscript.

